# Improving genome quality through artificial truncating purifying selection using heat shock: case of carps

**DOI:** 10.1101/2023.01.20.524958

**Authors:** Aleksandr Kuzmin, Vinogradov Evgenii, Dmitrii Balashov, Elena Kozenkova, Dmitrii Iliushchenko, Alina G. Mikhailova, Nataliya Patlay, Zakharenko Ekaterina, Kozenkov Ivan, Ptashnik Ivan, Valeriia Timonina, Evgenii Tretiakov, Yan Galimov, Christoph Haag, Bogdan Efimenko, Konstantin Gunbin, Igor Mizgirev, Nikolai Mugue, Konstantin Popadin

## Abstract

The process of domestication is associated with decrease in effective population size, which in turn leads to accumulation of slightly-deleterious mutations due to genetic drift. To maintain genome quality at a high level, we propose to use a stress-induced strong purifying selection, which based on negative epistasis, can effectively eliminate organisms with an excess of deleterious variants. Here, to identify stress factors, which interact with the effect of deleterious mutations we performed a proof-of-principle experiment with several regimes of a heat shock. We observed that fitness of mutated versus wild-type carp lines drops stronger after heat shock, which is a signature of a negative epistasis. Although the observed trend is promising, the effect of the epistasis is weak and unstable from family to family. Thus, more deep tuning of heat shock regimes is needed to uncover the most efficient combination of factors (absolute temperature, duration, stage of the embryo development) aggravating the burden of deleterious mutations and thus exposing them to the selection.

## Introduction

The process of domestication is associated with a degradation of genomes, characterized by an increase in the number, frequency, and severity of deleterious variants [1,2]. This degradation is primarily caused by a decrease in effective population size (Ne), resulting from bottlenecks, increased inbreeding, and the hitchhiking of deleterious variants linked to alleles under artificial selection. This genomic signature of domestication is universal for both plants and animals [3–5], including aquaculture species [6]. Thus, it is crucial to implement strategies to maintain a high level of genome quality in domesticated species, despite their relaxed purifying selection due to low Ne.

Deleterious variants, which are present in a population of domesticated species, individually have a low selective coefficient and therefore do not strongly impact fitness. However, the accumulation of hundreds or thousands of these variants can significantly affect an organism’s fitness in terms of health, fertility, and susceptibility to disease. Artificial selection against such variants is challenging to implement because it is hard to mimic natural selection that acts on inclusive fitness, which is a complex and nonlinear function of the genome and environment. Here we hypothesize that strong and universal stress can help artificially eliminate organisms with a high genome-wide burden of deleterious variants, thus compensating the relaxed natural selection in the domesticated populations. The logic behind this hypothesis is based on the negative epistasis. It has been shown that the majority of deleterious variants interact with each other through negative epistasis - i.e. mutually reinforcing each other’s deleterious effects, when the fitness effects is decreased relative to independence [7–9]. An increase in the strength of the negative epistasis between slightly deleterious variants, leading to more effective purifying selection can be induced by the presence of a severely deleterious genetic variant, “genetic handicap” [10]. Also, harsh or highly competitive environment can increase the strength of the negative epistasis between deleterious variants [11], thus increasing efficiency of the purifying selection. Thus by exposing organisms to environmental stress of the broad effect, we can intensify the negative interaction between deleterious variants, making selection more efficient.

What is the universal stress of broad effect? Here we define it as environmental factors affecting many unconditionally deleterious variants, i.e. variants which are deleterious across alternative environments because they affect essential functions of the organism [12]. Choosing among factors that are easy to implement for different model and nonmodel species, that are not mutagenic and that affect a broad spectrum of genes, we focused on a heat shock. First of all, it is well known that heat shock proteins buffer numerous slightly-deleterious variants segregating in genes-clients [13–16]. Prolonged heat shock is expected to exhaust the compensatory role of heat shock proteins (HSPs), revealing the previously hidden burden of slightly deleterious variants [17] and thus allowing selection to eliminate them. Second, heat shock can reveal previously hidden harmful variations not only among the hsp-clients (proteins that are normally assisted by heat shock proteins), but also in all other proteins. This is because an increase in temperature causes less stable, suboptimally folded structures with a higher abundance of harmful variations to become more prevalent. As a result, heat shock can be a widespread stressor that aggravates the effects of many cryptic deleterious variants.

For this experiment, carp - a freshwater fish with high fertility, external fertilization, and easy artificial fertilization was chosen as the model species. Heat shock was applied during the pharyngyla stage, when evolutionary old, haploinsufficient, and highly constrained genes are highly expressed [18]. We assume, that heat shock at this critical stage can be the most effective in the elimination of a burden of deleterious variants. Wild-type and mutated lines of carp were used to run a proof-of-principle experiment (Figure 2). We used diploid strain of the common carp (*Cyprinus carpio*) to facilitate mutagenic and heat shock experiments. If heat shock exacerbates the effect of deleterious variants (i.e. if there is a negative epistasis between the burden of deleterious variants and heat shock), it is expected to observe a more detrimental effect of heat shock on the mutated lines than the wild-type lines. If successful, the proof-of-principle experiment can be followed up with a purifying experiment, where the naturally existing variation in the burden of deleterious variants among siblings will be exposed to stress-induced purifying selection.

**Figure 1.**
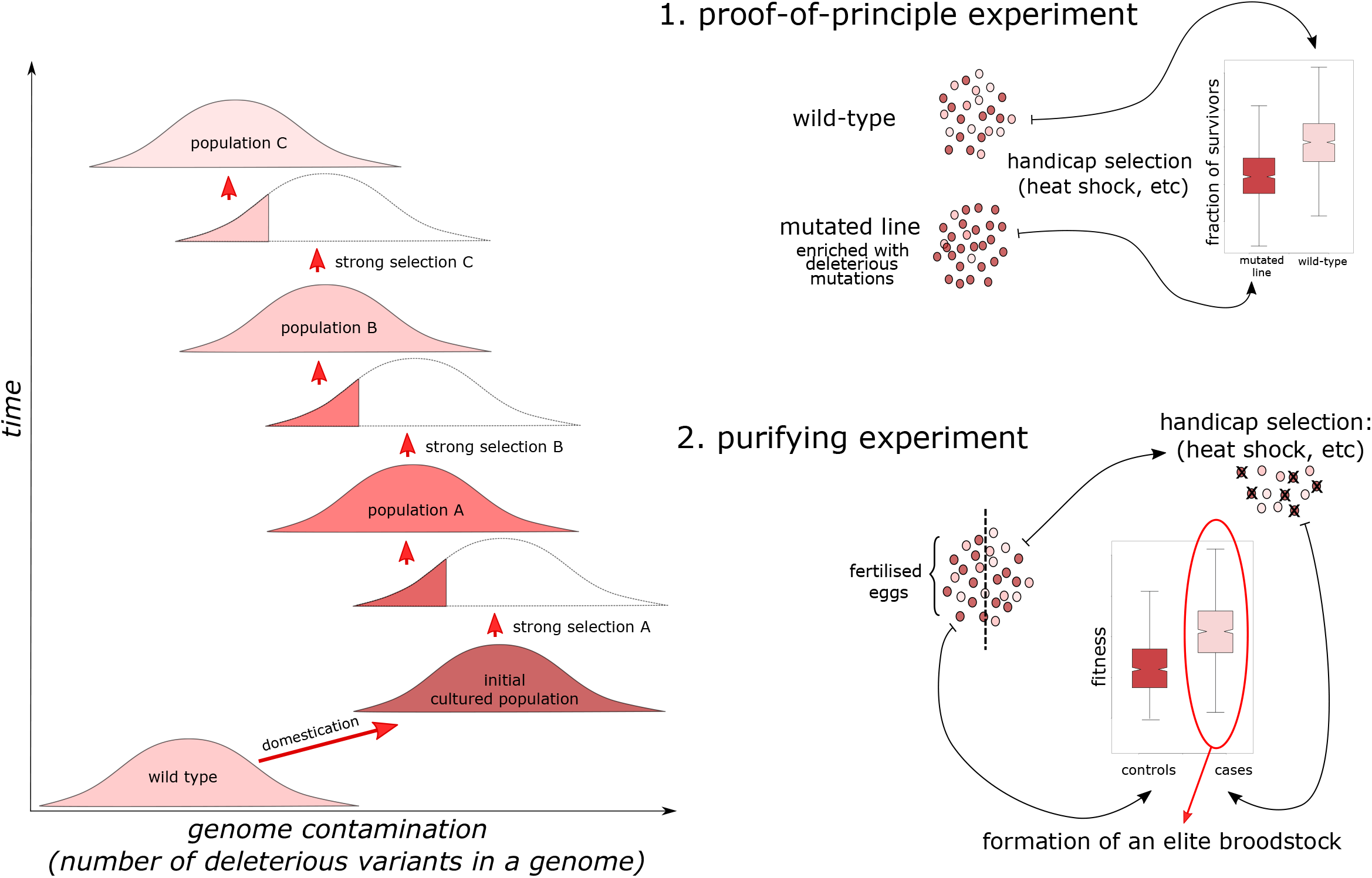
Left panel: Domestication is associated with genome degradation due to low effective population size. We propose that several rounds of strong purifying selection can improve genome quality back to the wild-type. Right upper panel: Proof-of-principle experiment. Right bottom panel: Purifying experiment

**Figure 2.**
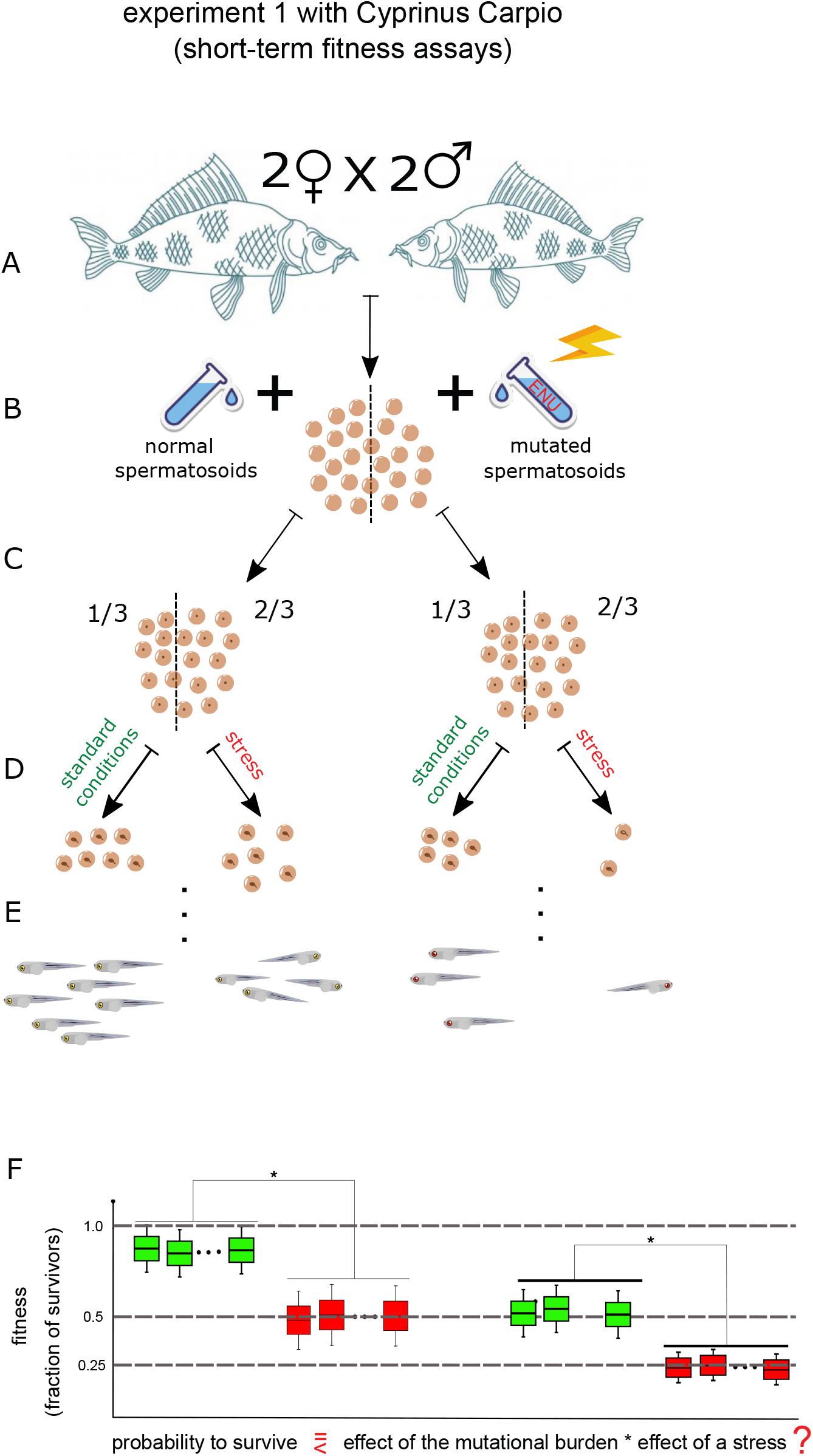
Scheme of a purifying experiment. A Obtaining of reproductive cells B ENU treatment of sperm C Sample preparation and fertilization of caviar D Heat shock of samples at 4 thermal modes E Incubation of larva up to a swimming stage F Expected survival probability is a function of mutational burden and stress

## Results

In the experiment, four families (two females mated with each of two males) were exposed to four different temperature regimes (controls at 21 degrees Celsius and cases at 38 degrees Celsius for 30, 40, and 50 minutes at the pharyngula stage) and three different mutagen regimes (no mutagen, 1.5, and 2 mM of ENU). This resulted in a total of 48 different combinations of families and regimes (4×4×3). Each of these combinations was repeated five times, resulting in a total of 240 petri dishes analyzed. Three primary, fitness-related traits were evaluated: the percentage of fertilized eggs, the percentage of eggs that hatched, and the percentage of properly swimming juveniles. The table with all results can be found as supplementary file 1.

All three primary traits showed a decrease following heat shock and following exposure to the mutagen (Figure 3, File S1). Among the three traits, the fraction of hatched eggs stood out as the most suitable for further analysis as it was the most stable, had a straightforward experimental setup, and had a clear biological interpretation.

**Figure 3.**
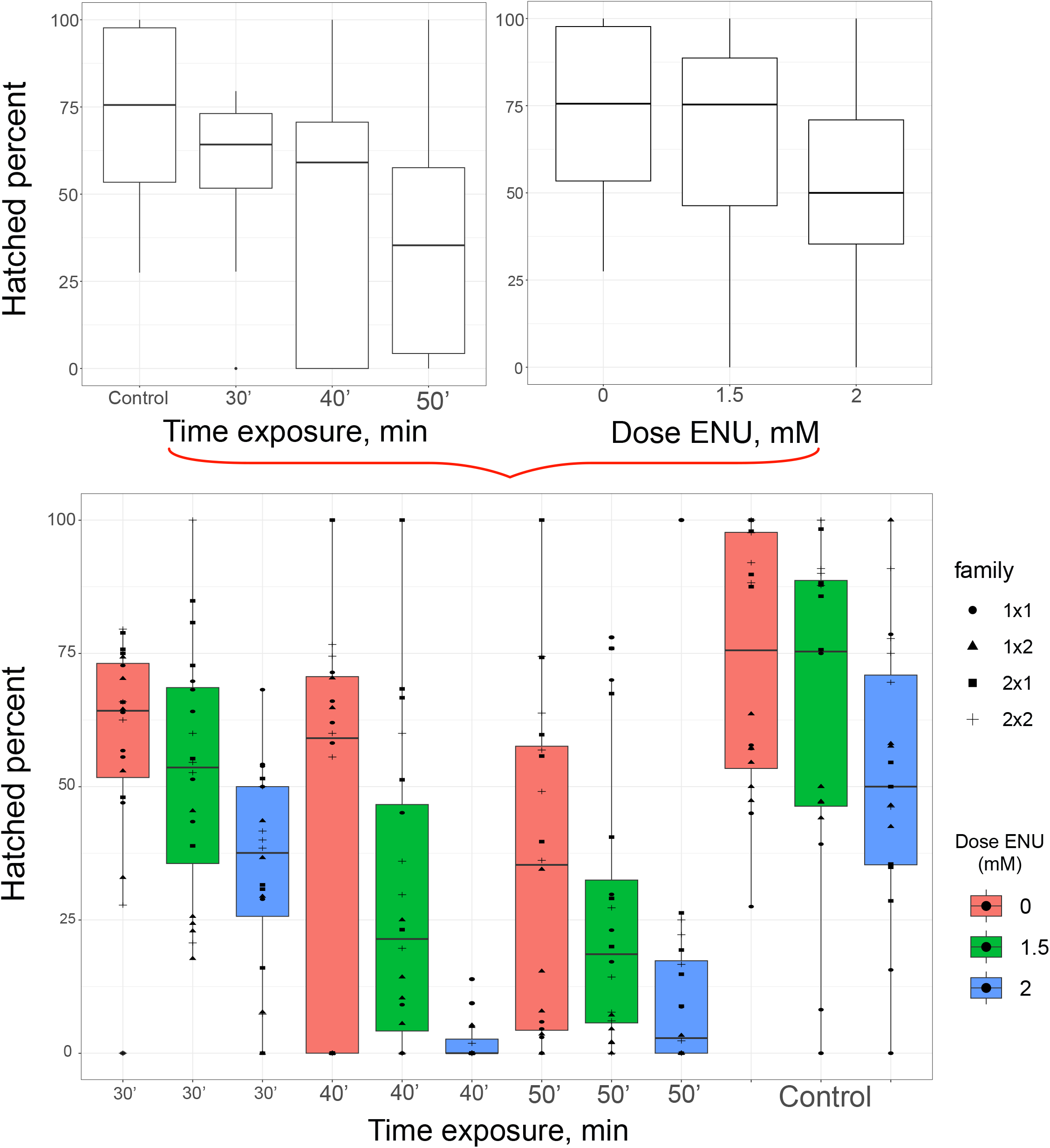
Upper panel: 2 boxplots shows correlation between hatched percent and (on the left) time exposure and (on the right) Dose ENU Bottom panel: boxplots shows synergistic effect of time exposure and mutagenes treatment. All boxplots are separated by heat shock mode (38° C for 30 min, 40 min, 50 min)

To gain insight into the effects of heat shock and mutagen on the probability of egg hatching, it is advantageous to focus on genetically similar eggs, i.e. on offspring from the same male and female. The outcome variable for each fertilized egg was coded as 1 if it hatched and 0 if it did not. The independent variables were MutagenConcentration (coded as 0, 1.5, or 2) and HeatShockDuration (coded as 0, 30, 40, or 50 minutes). This analysis was performed separately for each of the four families (see results in the table 1).

**Table 1.** results of the logistic multiple regression, describing the binary dependent variable HatchedOrNot as a function of two independent variables HeatShockDuration and MutagenConcentration. All parameters were scaled, which allowed to build regression through zero (intercept equal zero) and compare effects of each trait with each other despite the different original units.

| family (female x male) | HeatShockDuration |  | MutagenConcentration |  |
| --- | --- | --- | --- | --- |
|  | coefficient | p-value | coefficient | p-value |
| 1x1 | -0.03241 | 0.108 | -0.10843 | 4.74e-08 |
| 1x2 | -0.40876 | <2e-16 | -0.20066 | <2e-16 |
| 2x1 | -0.23299 | <2e-16 | -0.34536 | <2e-16 |
| 2x2 | -0.42811 | <2e-16 | -0.34538 | <2e-16 |

We observed a strong and significant negative correlation between hatching and both independent variables, HeatShockDuration and MutagenConcentration, in three out of the four families studied (Table 1, Figure 3). However, the 1×1 family did not show an expected correlation between hatching and HeatShockDuration. Thus, we removed this family from further analysis, where we analyze an interaction between two independent variables.

The similarity of coefficients for HeatShockDuration and MutagenConcentration, as shown in Table 1, across all families except 1×1, suggests equivalent effects on hatching probability. This indicates that the experimental settings (such as the temperature and duration, as well as mutagen concentration) were well-tuned in the initial phase of the experiment (see Methods).

Next, in the three families we analysed a potential interaction between HeatShockDuration and MutagenConcentration using the same logistic multiple regression approach. Results are in the Table 2.

**Table 2.** results of the logistic multiple regression, describing the binary dependent variable HatchedOrNot as a function of two independent variables HeatShockDuration and MutagenConcentration and their interaction. All parameters were scaled, which allowed to build regression through zero (intercept equal zero) and compare effects of each trait with each other despite the different original units.

| family:<br>female<br>x<br>male | HeatShockDuration |  | MutagenConcentration |  | HeatShockDuration<br>x<br>MutagenConcentration |  |
| --- | --- | --- | --- | --- | --- | --- |
|  | coefficient | p-value | coefficient | p-value | coefficient | p-value |
| **1x2** | -0.40945 | <2e-16 | -0.19887 | <2e-16 | -0.01423 | 0.516 |
| **2x1** | -0.235063 | <2e-16 | -0.346064 | <2e-16 | -0.009832 | 0.591 |
| **2x2** | -0.45045 | <2e-16 | -0.34252 | <2e-16 | -0.11935 | 4.88e-10 |
| **1x2, 2x1, 2x2** | -0.34488 | <2e-16 | -0.29858 | <2e-16 | -0.03277 | 0.00355 |

We observed a significant negative interaction between the HeatShockDuration and MutagenConcentration in one family (2×2) while interaction was not significant in two others (1×2, 2×1). The combination of all three families together (1×2, 2×1, 2×2) still demonstrated this negative interaction, although it was weaker (coefficient of interaction −0.03277, p-value 0.00355). When we included into the last model (1×2, 2×1, 2×2) dummy variables coding the families (FemaleOne and MaleOne), the negative interaction remained significant (coefficient −0.04374, p-value 6.20e-05).

Altogether we demonstrated that negative interaction between HeatShockDuration and MutagenConcentration is a plausible way of interaction between two analyzed deleterious factors.

To demonstrate better that the heat shock decreases fitness stronger in mutated versus wild-type lines, we performed more focused analysis, where family and mutagen concentration were the same and only the heat shock was different (Table 3). For each combination of family and mutagen concentration we estimated the fraction of hatched under heat shock (Hatched cases, “Hca”/All cases, “Aca”) and the fraction of hatched without heat shock (Hatched controls, “Hco” / All controls, “Aco”) and afterwards we got a ratio of these two fractions (Hca / Aca) / (Hco / Aco) which shows the relative decrease in the fraction of hatched due to heat shock. According to the NULL hypothesis of no epistasis the relative decrease in the fraction of hatched should not depend on mutagen (Enu), however we can see by eye that the relative decrease is becoming more pronounced within each family starting from wild-type (Enu = 0), to mutated (Enu = 1.5 or 2). For example, in family 2×1 under heat shock during 50 minutes the decrease in the fraction of hatched (as compared to the heat shock controls of the same family) was 0.699 for 0 ENU, 0.523 for 1.5 ENU and 0.395 for 2 ENU. This is a hallmark of negative epistasis, which shows that relative fitness due to heat shock (relative fraction of hatched) drops in mutated lines.

**Table 3.**
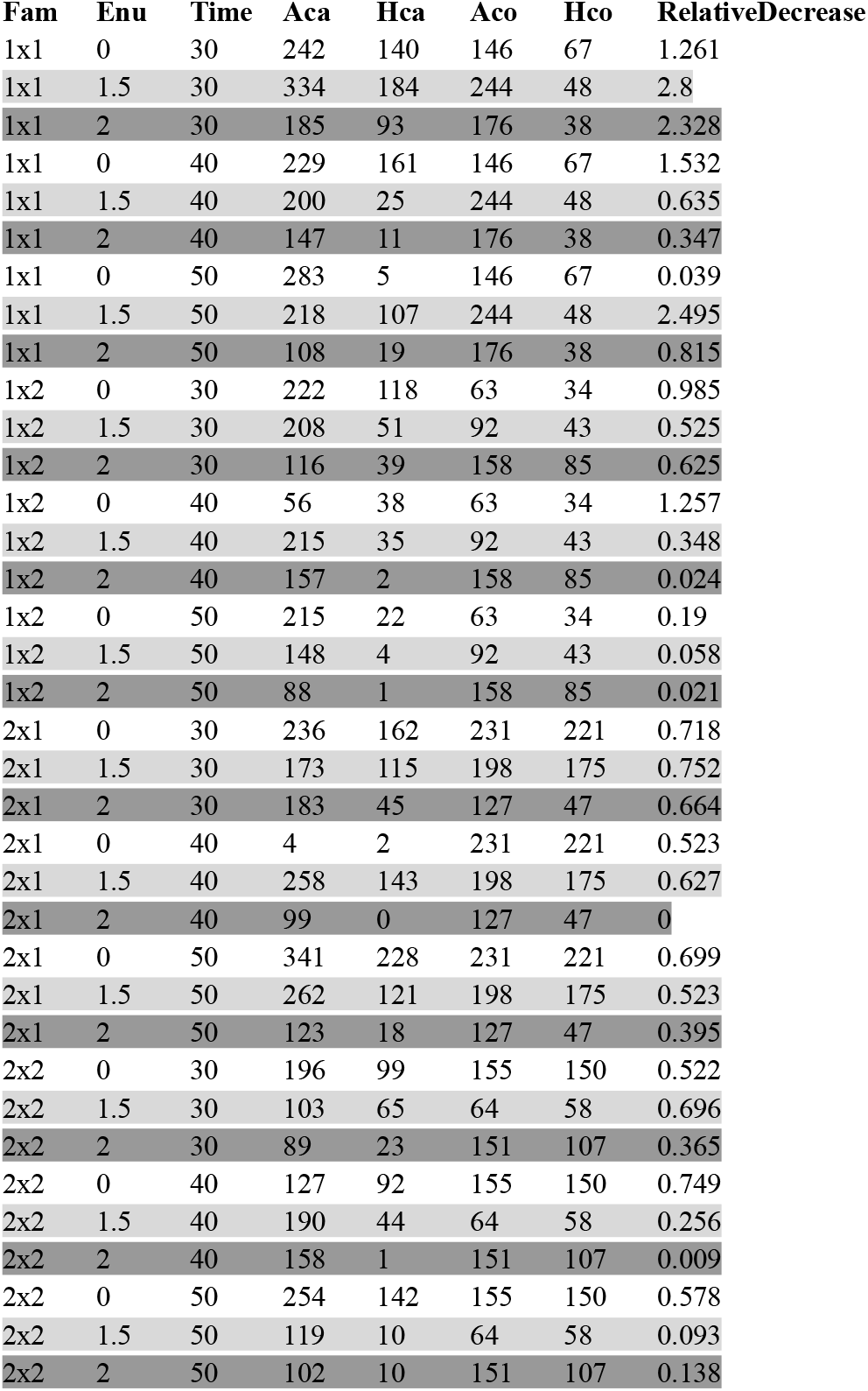
Family - and heat shock specific analysis, showing an increased effect of heat shock in mutated (dark grey and ligh grey phones) versus wild-type lines (white phone). Fam - family; Enu - concentration of ENU mutagen (mM), Time - time of the heat shock (minuts); Aca - All cases, total number of fertilized eggs in a heat shock affected groupt; Hca - hatched cases, number of hatched eggs in a heat shock affected group; Aco - All controls, number of fertilized eggs in the control group; Hco - Hatched controls, the number of hatched eggs in the control group.

Statistical analysis showed that after exclusion of family 1×1 (which demonstrates noisy and not expected increase in the percentage of hatched after the heat shock, see table 3) there is a marginally significant trend towards a reduction in the relative fraction of hatched eggs between 0 and 1.5 ENU (p = 0.074, paired two-sided Mann-Whitney U test, N=9+9), 0 and 2 ENU (p = 0.0040, paired two-sided Mann-Whitney U test, N=9+9) as well as 1.5 and 2 ENU (p = 0.055, paired two-sided Mann-Whitney U test, N=9+9).

## Discussion

In the current study, we planned to setup a stress-induced truncating purifying selection, which can reduce the burden of deleterious mutations in the genomes of domesticated species. This type of methodology might be implemented into cultivating fish populations to restore and maintain the high quality of the gene pool, which will contribute to increasing the economic value of fish farms.

To identify stress factors, which interact with the effect of harmful mutations we performed a proof-of-principle experiment. We observed that fitness (fraction of hatched fertilized embryos) of mutated versus wild-type lines drops stronger after heat shock, which is a signature of a negative epistasis. Although the observed trend is promising, the effect of the epistasis is weak and unstable from family to family. Thus, more deep tuning of heat shock regimes is needed to uncover the best (the most efficient) combination of factors (absolute temperature, duration, stage of the embryo development) aggravating the burden of deleterious mutations and thus exposing them to the selection.

## Materials and methods

### Add

ENU, sodium acetate, Hanks salts ect. chloroform was purchased from Reachim(Moscow, Russia). All chemicals were of analytical grade and used without further purification

The fertilization stage precedes the use of temperature shock. On pharyngula period of the common carp embryonic stages when organism reach formation of optic capsule and tail vesicle contraction commenced, and the embryo gradually moved with differentiation of most parts and a visible eye lens becoming more differentiated, then we could detect this “eye stage”(https://www.iasj.net/iasj/download/54e0817112cea846). When eggs reach “eye stage”, we shock them by LD50(t°C+exposure time)

Temperature is one of the main factors affecting the development of fish embryonic development. There is a known temperature range necessary for stimulation of sexual development, incubation of eggs and survival of young. This interval ranges from 19 to 30°C with an optimum around 23°C.

The stage of fertilization precedes the application of temperature shock. In the pharyngeal period of the embryonic stages of carp, the body forms visual capsules and the contraction of the tail vesicle begins, and the embryo gradually moves towards differentiation of most parts, in connection with which we can clearly distinguish the appearance of the lens of the eye - the “eye stage”(https://www.iasj.net/iasj/download/54e0817112cea846). It is at this stage that we apply heat shock.

According to the theory of the Hourglass model, we act on the most fragile stage of embryogenesis - the pharyngula, when multiple molecular networks are most connected between organ modules. In this way, we influence morphological divergence, because it is at this stage that most of the tissue laying takes place (https://doi.org/10.1242/dev.107318).

### Protocol

Before the final experiment we did 3 short-term experiments. In experiment 1 we tried to find LD50 for temperature, experiment 2 allowed us to determine LD50 for exposure time, also we added a mutagen to look at how they synergize with each other. Experiment 2 showed insufficient exposure to induced mutagen and we resorted to experiment 3, in which we selected the necessary LD50 concentration of ENU.

Finally, when we selected LD50 for temperature parameters(around 38 ° C, 40min) and LD50 for mutagen concentration(between 1.5 and 2.0 mM) we started the final, 4th experiment. During the experiment 4, the following parameters are taken into account: ENU concentrations (1.5 and 2.0 mM) Exposure time: 30, 40 and 50 minutes Shock temperature: 38.0 - 38.5 ° C.

#### Experiment 1

To create heat shock conditions, we used a short-term increase in ambient temperature (LD50) with different exposure modes in order to search for a half-year dose of elimination of 50% of individuals (eggs). Based on the literature, we have introduced 4 thermal modes: 30, 32, 34 and 36 ° C., as well as exposure time: 10, 15, 20, 25 minutes. In total, the study was conducted on 67 petri dishes, each of which contained 80-100 mg of caviar.

According to the results of the pilot experiment, we did not see a significant effect of induced heat shock (i.e., the death of eggs), as a result of which we developed two strategies for determining LD50 on these objects: (i) a stronger increase in temperature regime (which is likely to negatively affect embryonic development), (ii) prolongation of exposure time along with a slight increase in the temperature regime.

#### Experiment 2

The first large experiment for the selection of parameters of temperature shock with induced mutagenesis. During the experiment, the fertilization of carp eggs was carried out with sperm diluted at an ENU concentration of 0.55 mM (https://doi.org/10.1371/journal.pone.0026475) in solutions of sodium acetate (17 mM) and Hanks salts (9.8 mg/mL). To select LD50 according to the temperature regime of different exposure times, the parameters from experiment 1 were taken as the basis: we increased the exposure time and temperature exposure in order to break through the homeostatic barrier of the eggs. 80-100 mg of carp caviar is added to 288 petri dishes with a spoon and a spray of 1.25 mL is added. After applying caviar and sperm, 10 ml of aerated water is added to each Petri dish and mixing is performed. After 10 minutes of exposure, the water in the cups is replaced with fresh water (in order to remove unfertilized eggs and mutagen residues), the cups are placed in a thermostat for 24 hours at a temperature of 17.5 ° C.

A day later, the number of fertilized and dead eggs is estimated, and a day later, the assessment of hatching and swimming is carried out.

Data on the treatment of sperm with different exposure times and the presence of mutagen concentrations are shown in the table below:

| № Petri dish | Temperature | Time exposure, min | Concentration of ENU, mM |
| --- | --- | --- | --- |
| 1-16 | Control | Control | Control |
| 17-32 | Control | Control | 0,55 |
| 33-48,<br>161-176 | 37 | 30 | Control |
| 49-64,177-192 | 37 | 60 | Control |
| 65-80,<br>193-208 | 37 | 30 | 0,55 |
| 81-96,<br>209-224 | 37 | 60 | 0,55 |
| 97-112,<br>225-240 | 40 | 30 | Control |
| 113-128,<br>241-256 | 40 | 60 | Control |
| 129-144,<br>257-272 | 40 | 30 | 0,55 |
| 145-160,<br>273-288 | 40 | 60 | 0,55 |

#### Experiment 3

To select LD50 by ENU concentration for the final experiment, a smaller-scale study was performed. During the experiment, carp eggs were fertilized with sperm diluted at various concentrations of ENU in solutions of sodium acetate (17 mM) and Hanks salts (9.8 mg/mL). 80-100 mg of carp caviar is added to petri dishes with a spoon and sperm is added in a volume of 1.25 mL. Different concentrations of sperm treatment with mutagen are given in the table below:

| № Petri dish | Concentration of ENU, mM |
| --- | --- |
| 1-6 | 0 |
| 7-12 | 1,0 |
| 13-18 | 2,5 |
| 19-24 | 5,0 |
| 25-30 | 10,0 |

After applying caviar and sperm, 10 ml of aerated water is added to each Petri dish and mixing is performed. After 10 minutes of exposure, the water in the cups is replaced with fresh water (in order to remove unfertilized eggs and mutagen residues), the cups are placed in a thermostat for 24 hours at a temperature of 17.5 ° C.

A day later, the number of fertilized and dead eggs is estimated, and a day later, the assessment of hatching and swimming is carried out.

#### Experiment 4

Based on the data obtained in experiments 1, 2 and 3 we’ve made a fork of LD50 ENU concentrations and LD50 heat shock for the final experiment from 1.5 to 2.0 mM and 38 ° C with 40 min time exposure. As part of the final experiment, caviar from two females (No. 20 and 21) and sperm from two males (No. 25 and 28) were used, respectively. Sperm breeding is shown in the table below:

| Concentration of ENU, mM | Carp male №25 | Carp male №28 |
| --- | --- | --- |
| 1,5 | 4,75 mL Hanks' Balanced Salt,<br>73 uL ENU solution,<br>1,25 mL carp sperm | 4,75 mL Hanks' Balanced Salt,<br>75 uL ENU solution,<br>1,25 mL carp sperm |
| 2,0 | 4,9 mL Hanks' Balanced Salt,<br>100 uL ENU solution,<br>1,25 mL carp sperm | 4,9 mL Hanks' Balanced Salt,<br>100 uL ENU solution,<br>1,25 mL carp sperm |

The total exposure time after dilution is 45 minutes.

All the data and scripts are available at GitHub repository: https://github.com/mitoclub/fish-strong-purifying-selection

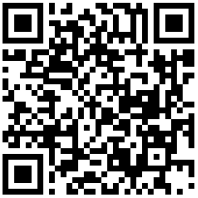

## Supporting information

https://github.com/mitoclub/fish-strong-purifying-selection/blob/master/figures/Supplementary_table_1.pdf

## Acknowledgments

AK and KP were supported by Ministry of Science and Higher Education of the Russian Federation (agreement no. 075-15-2021-1084) for design of the experiment, performance of the experiment and analysis of the results.

Branch for the freshwater fisheries of the Federal State Budget Scientific Institution “Russian Federal Research Institute of Fisheries and oceanography” Laboratory of genetics and fish breeding. Leading researcher

## Supplementary

table 1

