## Supplementary material for "Improving genome quality through artificial truncating purifying selection using heat shock: case of carps": https://github.com/mitoclub/fish-strong-purifying-selection/blob/master/figures/Supplementary_table_1.pdf: Supplementary_table_1.pdf

| petri | family | mode | mode_scr | temperature | time | enu | dead | alive | total | fertilization_per | hatched | hatched_per | swimming | freaks | swim_per | freaks_per | food | food_per |
|---|---|---|---|---|---|---|---|---|---|---|---|---|---|---|---|---|---|---|
| 1 | 1x1 | control | ctrl | 0 | 0 | 0 | 9 | 45 | 54 | 83,33 | 26 | 57,78 | 23 | 3 | 88,46 | 11,54 | | |
| 2 | 1x1 | control | ctrl | 0 | 0 | 0 | 4 | 20 | 24 | 83,33 | 9 | 45 | 5 | 4 | 55,56 | 44,44 | | |
| 3 | 1x1 | control | ctrl | 0 | 0 | 0 | 4 | 40 | 44 | 90,91 | 11 | 27,5 | 10 | 0 | 100 | 0 | | |
| 4 | 1x1 | control | ctrl | 0 | 0 | 0 | 2 | 20 | 22 | 90,91 | 9 | 45 | 8 | 1 | 88,89 | 11,11 | | |
| 5 | 1x1 | control | ctrl | 0 | 0 | 0 | 5 | 21 | 26 | 80,77 | 12 | 57,14 | 11 | 1 | 91,67 | 8,33 | | |
| 6 | 1x2 | control | ctrl | 0 | 0 | 0 | 2 | 14 | 16 | 87,5 | 8 | 57,14 | 4 | 4 | 50 | 50 | | |
| 7 | 1x2 | control | ctrl | 0 | 0 | 0 | 5 | 11 | 16 | 68,75 | 6 | 54,55 | 3 | 3 | 50 | 50 | | |
| 8 | 1x2 | control | ctrl | 0 | 0 | 0 | 4 | 8 | 12 | 66,67 | 4 | 50 | 2 | 2 | 50 | 50 | | |
| 9 | 1x2 | control | ctrl | 0 | 0 | 0 | 4 | 11 | 15 | 73,33 | 7 | 63,64 | 4 | 3 | 57,14 | 42,86 | | |
| 10 | 1x2 | control | ctrl | 0 | 0 | 0 | 1 | 19 | 20 | 95 | 9 | 47,37 | 5 | 4 | 55,56 | 44,44 | | |
| 11 | 2x1 | control | ctrl | 0 | 0 | 0 | 0 | 32 | 32 | 100 | 28 | 87,5 | 28 | 0 | 100 | 0 | | |
| 12 | 2x1 | control | ctrl | 0 | 0 | 0 | 0 | 29 | 29 | 100 | 33 | 113,79 | 29 | 1 | 96,67 | 3,33 | | |
| 13 | 2x1 | control | ctrl | 0 | 0 | 0 | 5 | 67 | 72 | 93,06 | 69 | 102,99 | 55 | 2 | 96,49 | 3,51 | | |
| 14 | 2x1 | control | ctrl | 0 | 0 | 0 | 0 | 48 | 48 | 100 | 47 | 97,92 | 39 | 0 | 100 | 0 | | |
| 15 | 2x1 | control | ctrl | 0 | 0 | 0 | 1 | 49 | 50 | 98 | 44 | 89,8 | 42 | 0 | 100 | 0 | | |
| 16 | 2x2 | control | ctrl | 0 | 0 | 0 | 1 | 42 | 43 | 97,67 | 41 | 97,62 | 38 | 2 | 95 | 5 | | |
| 17 | 2x2 | control | ctrl | 0 | 0 | 0 | 0 | 17 | 17 | 100 | 15 | 88,24 | 15 | 0 | 100 | 0 | | |
| 18 | 2x2 | control | ctrl | 0 | 0 | 0 | 4 | 40 | 44 | 90,91 | 40 | 100 | 37 | 3 | 92,5 | 7,5 | | |
| 19 | 2x2 | control | ctrl | 0 | 0 | 0 | 1 | 29 | 30 | 96,67 | 31 | 106,9 | 28 | 3 | 90,32 | 9,68 | | |
| 20 | 2x2 | control | ctrl | 0 | 0 | 0 | 2 | 25 | 27 | 92,59 | 23 | 92 | 19 | 3 | 86,36 | 13,64 | | |
| 21 | 1x1 | control_enu_1 | ctrl_e1 | 0 | 0 | 1 | 15 | 49 | 64 | 76,56 | 4 | 8,16 | 4 | 0 | 100 | 0 | | |
| 22 | 1x1 | control_enu_1 | ctrl_e1 | 0 | 0 | 1 | 16 | 69 | 85 | 81,18 | 0 | 0 | 0 | 0 | 0 | 0 | | |
| 23 | 1x1 | control_enu_1 | ctrl_e1 | 0 | 0 | 1 | 12 | 43 | 55 | 78,18 | 0 | 0 | 0 | 0 | 0 | 0 | | |
| 24 | 1x1 | control_enu_1 | ctrl_e1 | 0 | 0 | 1 | 27 | 51 | 78 | 65,38 | 20 | 39,22 | 17 | 3 | 85 | 15 | | |
| 25 | 1x1 | control_enu_1 | ctrl_e1 | 0 | 0 | 1 | 9 | 32 | 41 | 78,05 | 24 | 75 | 16 | 2 | 88,89 | 11,11 | | |
| 26 | 1x2 | control_enu_1 | ctrl_e1 | 0 | 0 | 1 | 16 | 36 | 52 | 69,23 | 17 | 47,22 | 7 | 10 | 41,18 | 58,82 | | |
| 27 | 1x2 | control_enu_1 | ctrl_e1 | 0 | 0 | 1 | 53 | 10 | 63 | 15,87 | 5 | 50 | 1 | 4 | 20 | 80 | | |
| 28 | 1x2 | control_enu_1 | ctrl_e1 | 0 | 0 | 1 | 31 | 34 | 65 | 52,31 | 15 | 44,12 | 11 | 4 | 73,33 | 26,67 | | |
| 29 | 1x2 | control_enu_1 | ctrl_e1 | 0 | 0 | 1 | 31 | 12 | 43 | 27,91 | 6 | 50 | 3 | 3 | 50 | 50 | | |
| 30 | 1x2 | control_enu_1 | ctrl_e1 | 0 | 0 | 1 | 63 | 34 | 97 | 35,05 | 16 | 47,06 | 12 | 5 | 70,59 | 29,41 | | |
| 31 | 2x1 | control_enu_1 | ctrl_e1 | 0 | 0 | 1 | 3 | 34 | 37 | 91,89 | 30 | 88,24 | 30 | 8 | 78,95 | 21,05 | | |
| 32 | 2x1 | control_enu_1 | ctrl_e1 | 0 | 0 | 1 | 9 | 37 | 46 | 80,43 | 28 | 75,68 | 15 | 13 | 53,57 | 46,43 | | |
| 33 | 2x1 | control_enu_1 | ctrl_e1 | 0 | 0 | 1 | 3 | 28 | 31 | 90,32 | 24 | 85,71 | 17 | 6 | 73,91 | 26,09 | | |
| 34 | 2x1 | control_enu_1 | ctrl_e1 | 0 | 0 | 1 | 2 | 74 | 76 | 97,37 | 65 | 87,84 | 43 | 22 | 66,15 | 33,85 | | |
| 35 | 2x1 | control_enu_1 | ctrl_e1 | 0 | 0 | 1 | 7 | 59 | 66 | 89,39 | 58 | 98,31 | 21 | 37 | 36,21 | 63,79 | | |
| 36 | 2x2 | control_enu_1 | ctrl_e1 | 0 | 0 | 1 | 3 | 13 | 16 | 81,25 | 13 | 100 | 2 | 12 | 14,29 | 85,71 | | |
| 37 | 2x2 | control_enu_1 | ctrl_e1 | 0 | 0 | 1 | 25 | 20 | 45 | 44,44 | 18 | 90 | 10 | 7 | 58,82 | 41,18 | | |
| 38 | 2x2 | control_enu_1 | ctrl_e1 | 0 | 0 | 1 | 7 | 57 | 64 | 89,06 | 50 | 87,72 | 22 | 30 | 42,31 | 57,69 | | |
| 39 | 2x2 | control_enu_1 | ctrl_e1 | 0 | 0 | 1 | 15 | 22 | 37 | 59,46 | 22 | 100 | 7 | 16 | 30,43 | 69,57 | | |
| 40 | 2x2 | control_enu_1 | ctrl_e1 | 0 | 0 | 1 | 22 | 44 | 66 | 66,67 | 40 | 90,91 | 20 | 20 | 50 | 50 | | |
| 41 | 1x1 | control_enu_2 | ctrl_e2 | 0 | 0 | 2 | 11 | 42 | 53 | 79,25 | 0 | 0 | 0 | 0 | 0 | 0 | | |
| 42 | 1x1 | control_enu_2 | ctrl_e2 | 0 | 0 | 2 | 25 | 52 | 77 | 67,53 | 0 | 0 | 0 | 0 | 0 | 0 | | |
| 43 | 1x1 | control_enu_2 | ctrl_e2 | 0 | 0 | 2 | 21 | 32 | 53 | 60,38 | 5 | 15,63 | 2 | 3 | 40 | 60 | | |
| 44 | 1x1 | control_enu_2 | ctrl_e2 | 0 | 0 | 2 | 31 | 22 | 53 | 41,51 | 11 | 50 | 7 | 4 | 63,64 | 36,36 | | |
| 45 | 1x1 | control_enu_2 | ctrl_e2 | 0 | 0 | 2 | 25 | 28 | 53 | 52,83 | 22 | 78,57 | 12 | 10 | 54,55 | 45,45 | | |
| 46 | 1x2 | control_enu_2 | ctrl_e2 | 0 | 0 | 2 | 34 | 31 | 65 | 47,69 | 18 | 58,06 | 14 | 4 | 77,78 | 22,22 | | |
| 47 | 1x2 | control_enu_2 | ctrl_e2 | 0 | 0 | 2 | 61 | 33 | 94 | 35,11 | 19 | 57,58 | 13 | 6 | 68,42 | 31,58 | | |
| 48 | 1x2 | control_enu_2 | ctrl_e2 | 0 | 0 | 2 | 20 | 10 | 30 | 33,33 | 11 | 110 | 6 | 5 | 54,55 | 45,45 | | |
| 49 | 1x2 | control_enu_2 | ctrl_e2 | 0 | 0 | 2 | 70 | 40 | 110 | 36,36 | 17 | 42,5 | 13 | 4 | 76,47 | 23,53 | | |
| 50 | 1x2 | control_enu_2 | ctrl_e2 | 0 | 0 | 2 | 40 | 43 | 83 | 51,81 | 20 | 46,51 | 15 | 5 | 75 | 25 | | |
| 51 | 2x1 | control_enu_2 | ctrl_e2 | 0 | 0 | 2 | 52 | 43 | 95 | 45,26 | 15 | 34,88 | 11 | 4 | 73,33 | 26,67 | | |

| | | | | | | | | | | | | | | | | |
|---|---|---|---|---|---|---|---|---|---|---|---|---|---|---|---|---|
| 52 | 2x1 | control_enu_2 | ctrl_e2 | 0 | 0 | 2 | 21 | 31 | 52 | 59,62 | 11 | 35,48 | 1 | 10 | 9,09 | 90,91 |
| 53 | 2x1 | control_enu_2 | ctrl_e2 | 0 | 0 | 2 | 6 | 14 | 20 | 70 | 7 | 50 | 3 | 3 | 50 | 50 |
| 54 | 2x1 | control_enu_2 | ctrl_e2 | 0 | 0 | 2 | 6 | 11 | 17 | 64,71 | 6 | 54,55 | 4 | 2 | 66,67 | 33,33 |
| 55 | 2x1 | control_enu_2 | ctrl_e2 | 0 | 0 | 2 | 14 | 28 | 42 | 66,67 | 8 | 28,57 | 3 | 4 | 42,86 | 57,14 |
| 56 | 2x2 | control_enu_2 | ctrl_e2 | 0 | 0 | 2 | 22 | 48 | 70 | 68,57 | 36 | 75 | 9 | 26 | 25,71 | 74,29 |
| 57 | 2x2 | control_enu_2 | ctrl_e2 | 0 | 0 | 2 | 42 | 46 | 88 | 52,27 | 32 | 69,57 | 11 | 21 | 34,38 | 65,63 |
| 58 | 2x2 | control_enu_2 | ctrl_e2 | 0 | 0 | 2 | 15 | 22 | 37 | 59,46 | 20 | 90,91 | 6 | 14 | 30 | 70 |
| 59 | 2x2 | control_enu_2 | ctrl_e2 | 0 | 0 | 2 | 31 | 26 | 57 | 45,61 | 12 | 46,15 | 5 | 7 | 41,67 | 58,33 |
| 60 | 2x2 | control_enu_2 | ctrl_e2 | 0 | 0 | 2 | 26 | 9 | 35 | 25,71 | 7 | 77,78 | 2 | 5 | 28,57 | 71,43 |
| 61 | 1x1 | 38_30_min_nat | 38_30_nat | 38 | 30 | 0 | 5 | 37 | 42 | 88,1 | 21 | 56,76 | 20 | 0 | 100 | 0 |
| 62 | 1x1 | 38_30_min_nat | 38_30_nat | 38 | 30 | 0 | 6 | 33 | 39 | 84,62 | 24 | 72,73 | 24 | 0 | 100 | 0 |
| 63 | 1x1 | 38_30_min_nat | 38_30_nat | 38 | 30 | 0 | 15 | 61 | 76 | 80,26 | 39 | 63,93 | 37 | 0 | 100 | 0 |
| 64 | 1x1 | 38_30_min_nat | 38_30_nat | 38 | 30 | 0 | 13 | 66 | 79 | 83,54 | 31 | 46,97 | 27 | 3 | 90 | 10 |
| 65 | 1x1 | 38_30_min_nat | 38_30_nat | 38 | 30 | 0 | 8 | 45 | 53 | 84,91 | 25 | 55,56 | 21 | 1 | 95,45 | 4,55 |
| 66 | 1x2 | 38_30_min_nat | 38_30_nat | 38 | 30 | 0 | 6 | 37 | 43 | 86,05 | 26 | 70,27 | 24 | 2 | 92,31 | 7,69 |
| 67 | 1x2 | 38_30_min_nat | 38_30_nat | 38 | 30 | 0 | 12 | 35 | 47 | 74,47 | 26 | 74,29 | 19 | 2 | 90,48 | 9,52 |
| 68 | 1x2 | 38_30_min_nat | 38_30_nat | 38 | 30 | 0 | 7 | 31 | 38 | 81,58 | 20 | 64,52 | 13 | 2 | 86,67 | 13,33 |
| 69 | 1x2 | 38_30_min_nat | 38_30_nat | 38 | 30 | 0 | 15 | 85 | 100 | 85 | 28 | 32,94 | 21 | 2 | 91,3 | 8,7 |
| 70 | 1x2 | 38_30_min_nat | 38_30_nat | 38 | 30 | 0 | 6 | 34 | 40 | 85 | 18 | 52,94 | 12 | 1 | 92,31 | 7,69 |
| 71 | 2x1 | 38_30_min_nat | 38_30_nat | 38 | 30 | 0 | 0 | 33 | 33 | 100 | 25 | 75,76 | 18 | 7 | 72 | 28 |
| 72 | 2x1 | 38_30_min_nat | 38_30_nat | 38 | 30 | 0 | 2 | 60 | 62 | 96,77 | 45 | 75 | 40 | 3 | 93,02 | 6,98 |
| 73 | 2x1 | 38_30_min_nat | 38_30_nat | 38 | 30 | 0 | 8 | 50 | 58 | 86,21 | 24 | 48 | 20 | 3 | 86,96 | 13,04 |
| 74 | 2x1 | 38_30_min_nat | 38_30_nat | 38 | 30 | 0 | 1 | 52 | 53 | 98,11 | 41 | 78,85 | 37 | 2 | 94,87 | 5,13 |
| 75 | 2x1 | 38_30_min_nat | 38_30_nat | 38 | 30 | 0 | 0 | 41 | 41 | 100 | 27 | 65,85 | 21 | 4 | 84 | 16 |
| 76 | 2x2 | 38_30_min_nat | 38_30_nat | 38 | 30 | 0 | 2 | 44 | 46 | 95,65 | 29 | 65,91 | 14 | 11 | 56 | 44 |
| 77 | 2x2 | 38_30_min_nat | 38_30_nat | 38 | 30 | 0 | 5 | 40 | 45 | 88,89 | 25 | 62,5 | 11 | 10 | 52,38 | 47,62 |
| 78 | 2x2 | 38_30_min_nat | 38_30_nat | 38 | 30 | 0 | 2 | 32 | 34 | 94,12 | 0 | 0 | 0 | 0 | 0 | 0 |
| 79 | 2x2 | 38_30_min_nat | 38_30_nat | 38 | 30 | 0 | 5 | 36 | 41 | 87,8 | 10 | 27,78 | 7 | 3 | 70 | 30 |
| 80 | 2x2 | 38_30_min_nat | 38_30_nat | 38 | 30 | 0 | 3 | 44 | 47 | 93,62 | 35 | 79,55 | 24 | 11 | 68,57 | 31,43 |
| 81 | 1x1 | 38_40_min_nat | 38_40_nat | 38 | 40 | 0 | 11 | 50 | 61 | 81,97 | 31 | 62 | 29 | 2 | 93,55 | 6,45 |
| 82 | 1x1 | 38_40_min_nat | 38_40_nat | 38 | 40 | 0 | 11 | 53 | 64 | 82,81 | 35 | 66,04 | 30 | 5 | 85,71 | 14,29 |
| 83 | 1x1 | 38_40_min_nat | 38_40_nat | 38 | 40 | 0 | 5 | 28 | 33 | 84,85 | 20 | 71,43 | 18 | 2 | 90 | 10 |
| 84 | 1x1 | 38_40_min_nat | 38_40_nat | 38 | 40 | 0 | 15 | 55 | 70 | 78,57 | 32 | 58,18 | 27 | 5 | 84,38 | 15,63 |
| 85 | 1x1 | 38_40_min_nat | 38_40_nat | 38 | 40 | 0 | 6 | 41 | 47 | 87,23 | 43 | 104,88 | 32 | 11 | 74,42 | 25,58 |
| 86 | 1x2 | 38_40_min_nat | 38_40_nat | 38 | 40 | 0 | 4 | 54 | 58 | 93,1 | 38 | 70,37 | 30 | 8 | 78,95 | 21,05 |
| 87 | 1x2 | 38_40_min_nat | 38_40_nat | 38 | 40 | 0 | 20 | 71 | 91 | 78,02 | 46 | 64,79 | 36 | 11 | 76,6 | 23,4 |
| 88 | 1x2 | 38_40_min_nat | 38_40_nat | 38 | 40 | 0 | 17 | 2 | 19 | 10,53 | 0 | 0 | 0 | 0 | 0 | 0 |
| 89 | 1x2 | 38_40_min_nat | 38_40_nat | 38 | 40 | 0 | 61 | 0 | 61 | 0 | 0 | 0 | 0 | 0 | 0 | 0 |
| 90 | 1x2 | 38_40_min_nat | 38_40_nat | 38 | 40 | 0 | 50 | 0 | 50 | 0 | 0 | 0 | 0 | 0 | 0 | 0 |
| 91 | 2x1 | 38_40_min_nat | 38_40_nat | 38 | 40 | 0 | 21 | 2 | 23 | 8,7 | 2 | 100 | 2 | 0 | 100 | 0 |
| 92 | 2x1 | 38_40_min_nat | 38_40_nat | 38 | 40 | 0 | 23 | 1 | 24 | 4,17 | 0 | 0 | 0 | 0 | 0 | 0 |
| 93 | 2x1 | 38_40_min_nat | 38_40_nat | 38 | 40 | 0 | 50 | 0 | 50 | 0 | 0 | 0 | 0 | 0 | 0 | 0 |
| 94 | 2x1 | 38_40_min_nat | 38_40_nat | 38 | 40 | 0 | 35 | 1 | 36 | 2,78 | 0 | 0 | 0 | 0 | 0 | 0 |
| 95 | 2x1 | 38_40_min_nat | 38_40_nat | 38 | 40 | 0 | 50 | 0 | 50 | 0 | 0 | 0 | 0 | 0 | 0 | 0 |
| 96 | 2x2 | 38_40_min_nat | 38_40_nat | 38 | 40 | 0 | 25 | 1 | 26 | 3,85 | 0 | 0 | 0 | 0 | 0 | 0 |
| 97 | 2x2 | 38_40_min_nat | 38_40_nat | 38 | 40 | 0 | 24 | 9 | 33 | 27,27 | 5 | 55,56 | 0 | 5 | 0 | 100 |
| 98 | 2x2 | 38_40_min_nat | 38_40_nat | 38 | 40 | 0 | 8 | 10 | 18 | 55,56 | 6 | 60 | 3 | 3 | 50 | 50 |
| 99 | 2x2 | 38_40_min_nat | 38_40_nat | 38 | 40 | 0 | 7 | 47 | 54 | 87,04 | 35 | 74,47 | 20 | 15 | 57,14 | 42,86 |
| 100 | 2x2 | 38_40_min_nat | 38_40_nat | 38 | 40 | 0 | 4 | 60 | 64 | 93,75 | 46 | 76,67 | 36 | 10 | 78,26 | 21,74 |
| 101 | 1x1 | 38_50_min_nat | 38_50_nat | 38 | 50 | 0 | 21 | 87 | 108 | 80,56 | 0 | 0 | 0 | 0 | 0 | 0 |
| 102 | 1x1 | 38_50_min_nat | 38_50_nat | 38 | 50 | 0 | 18 | 73 | 91 | 80,22 | 0 | 0 | 0 | 0 | 0 | 0 |
| 103 | 1x1 | 38_50_min_nat | 38_50_nat | 38 | 50 | 0 | 11 | 34 | 45 | 75,56 | 2 | 5,88 | 1 | 1 | 50 | 50 |

| | | | | | | | | | | | | | | | | |
|---|---|---|---|---|---|---|---|---|---|---|---|---|---|---|---|---|
| 104 | 1x1 | 38_50_min_nat | 38_50_nat | 38 | 50 | 0 | 11 | 67 | 78 | 85,9 | 2 | 2,99 | 0 | 2 | 0 | 100 |
| 105 | 1x1 | 38_50_min_nat | 38_50_nat | 38 | 50 | 0 | 7 | 22 | 29 | 75,86 | 1 | 4,55 | 1 | 0 | 100 | 0 |
| 106 | 1x2 | 38_50_min_nat | 38_50_nat | 38 | 50 | 0 | 11 | 29 | 40 | 72,5 | 10 | 34,48 | 5 | 5 | 50 | 50 |
| 107 | 1x2 | 38_50_min_nat | 38_50_nat | 38 | 50 | 0 | 7 | 52 | 59 | 88,14 | 8 | 15,38 | 6 | 2 | 75 | 25 |
| 108 | 1x2 | 38_50_min_nat | 38_50_nat | 38 | 50 | 0 | 6 | 28 | 34 | 82,35 | 1 | 3,57 | 1 | 0 | 100 | 0 |
| 109 | 1x2 | 38_50_min_nat | 38_50_nat | 38 | 50 | 0 | 15 | 68 | 83 | 81,93 | 0 | 0 | 0 | 0 | 0 | 0 |
| 110 | 1x2 | 38_50_min_nat | 38_50_nat | 38 | 50 | 0 | 8 | 38 | 46 | 82,61 | 3 | 7,89 | 2 | 1 | 66,67 | 33,33 |
| 111 | 2x1 | 38_50_min_nat | 38_50_nat | 38 | 50 | 0 | 0 | 74 | 74 | 100 | 74 | 100 | 66 | 8 | 89,19 | 10,81 |
| 112 | 2x1 | 38_50_min_nat | 38_50_nat | 38 | 50 | 0 | 3 | 66 | 69 | 95,65 | 49 | 74,24 | 40 | 6 | 86,96 | 13,04 |
| 113 | 2x1 | 38_50_min_nat | 38_50_nat | 38 | 50 | 0 | 1 | 63 | 64 | 98,44 | 25 | 39,68 | 20 | 2 | 90,91 | 9,09 |
| 114 | 2x1 | 38_50_min_nat | 38_50_nat | 38 | 50 | 0 | 1 | 61 | 62 | 98,39 | 34 | 55,74 | 26 | 8 | 76,47 | 23,53 |
| 115 | 2x1 | 38_50_min_nat | 38_50_nat | 38 | 50 | 0 | 3 | 77 | 80 | 96,25 | 46 | 59,74 | 36 | 10 | 78,26 | 21,74 |
| 116 | 2x2 | 38_50_min_nat | 38_50_nat | 38 | 50 | 0 | 4 | 55 | 59 | 93,22 | 27 | 49,09 | 18 | 8 | 69,23 | 30,77 |
| 117 | 2x2 | 38_50_min_nat | 38_50_nat | 38 | 50 | 0 | 5 | 58 | 63 | 92,06 | 37 | 63,79 | 23 | 10 | 69,7 | 30,3 |
| 118 | 2x2 | 38_50_min_nat | 38_50_nat | 38 | 50 | 0 | 0 | 43 | 43 | 100 | 32 | 74,42 | 21 | 9 | 70 | 30 |
| 119 | 2x2 | 38_50_min_nat | 38_50_nat | 38 | 50 | 0 | 3 | 47 | 50 | 94 | 17 | 36,17 | 10 | 6 | 62,5 | 37,5 |
| 120 | 2x2 | 38_50_min_nat | 38_50_nat | 38 | 50 | 0 | 2 | 51 | 53 | 96,23 | 29 | 56,86 | 17 | 8 | 68 | 32 |
| 121 | 1x1 | 38_30_min_EN | 38_30_en1 | 38 | 30 | 1 | 7 | 43 | 50 | 86 | 30 | 69,77 | 24 | 6 | 80 | 20 |
| 122 | 1x1 | 38_30_min_EN | 38_30_en1 | 38 | 30 | 1 | 16 | 109 | 125 | 87,2 | 56 | 51,38 | 26 | 1 | 96,3 | 3,7 |
| 123 | 1x1 | 38_30_min_EN | 38_30_en1 | 38 | 30 | 1 | 19 | 99 | 118 | 83,9 | 43 | 43,43 | 36 | 7 | 83,72 | 16,28 |
| 124 | 1x1 | 38_30_min_EN | 38_30_en1 | 38 | 30 | 1 | 10 | 39 | 49 | 79,59 | 25 | 64,1 | 17 | 8 | 68 | 32 |
| 125 | 1x1 | 38_30_min_EN | 38_30_en1 | 38 | 30 | 1 | 5 | 44 | 49 | 89,8 | 30 | 68,18 | 23 | 6 | 79,31 | 20,69 |
| 126 | 1x2 | 38_30_min_EN | 38_30_en1 | 38 | 30 | 1 | 21 | 39 | 60 | 65 | 10 | 25,64 | 7 | 3 | 70 | 30 |
| 127 | 1x2 | 38_30_min_EN | 38_30_en1 | 38 | 30 | 1 | 44 | 62 | 106 | 58,49 | 11 | 17,74 | 8 | 2 | 80 | 20 |
| 128 | 1x2 | 38_30_min_EN | 38_30_en1 | 38 | 30 | 1 | 22 | 37 | 59 | 62,71 | 9 | 24,32 | 3 | 4 | 42,86 | 57,14 |
| 129 | 1x2 | 38_30_min_EN | 38_30_en1 | 38 | 30 | 1 | 18 | 22 | 40 | 55 | 10 | 45,45 | 4 | 6 | 40 | 60 |
| 130 | 1x2 | 38_30_min_EN | 38_30_en1 | 38 | 30 | 1 | 18 | 48 | 66 | 72,73 | 11 | 22,92 | 8 | 3 | 72,73 | 27,27 |
| 131 | 2x1 | 38_30_min_EN | 38_30_en1 | 38 | 30 | 1 | 5 | 33 | 38 | 86,84 | 24 | 72,73 | 12 | 9 | 57,14 | 42,86 |
| 132 | 2x1 | 38_30_min_EN | 38_30_en1 | 38 | 30 | 1 | 15 | 66 | 81 | 81,48 | 56 | 84,85 | 29 | 23 | 55,77 | 44,23 |
| 133 | 2x1 | 38_30_min_EN | 38_30_en1 | 38 | 30 | 1 | 7 | 26 | 33 | 78,79 | 21 | 80,77 | 9 | 12 | 42,86 | 57,14 |
| 134 | 2x1 | 38_30_min_EN | 38_30_en1 | 38 | 30 | 1 | 24 | 36 | 60 | 60 | 14 | 38,89 | 13 | 1 | 92,86 | 7,14 |
| 135 | 2x1 | 38_30_min_EN | 38_30_en1 | 38 | 30 | 1 | 15 | 38 | 53 | 71,7 | 21 | 55,26 | 14 | 6 | 70 | 30 |
| 136 | 2x2 | 38_30_min_EN | 38_30_en1 | 38 | 30 | 1 | 8 | 29 | 37 | 78,38 | 6 | 20,69 | 3 | 3 | 50 | 50 |
| 137 | 2x2 | 38_30_min_EN | 38_30_en1 | 38 | 30 | 1 | 18 | 31 | 49 | 63,27 | 38 | 122,58 | 31 | 7 | 81,58 | 18,42 |
| 138 | 2x2 | 38_30_min_EN | 38_30_en1 | 38 | 30 | 1 | 10 | 11 | 21 | 52,38 | 6 | 54,55 | 5 | 1 | 83,33 | 16,67 |
| 139 | 2x2 | 38_30_min_EN | 38_30_en1 | 38 | 30 | 1 | 24 | 19 | 43 | 44,19 | 10 | 52,63 | 24 | 5 | 82,76 | 17,24 |
| 140 | 2x2 | 38_30_min_EN | 38_30_en1 | 38 | 30 | 1 | 26 | 25 | 51 | 49,02 | 15 | 60 | 5 | 0 | 100 | 0 |
| 141 | 1x1 | 38_40_min_EN | 38_40_en1 | 38 | 40 | 1 | 11 | 47 | 58 | 81,03 | 0 | 0 | 0 | 0 | 0 | 0 |
| 142 | 1x1 | 38_40_min_EN | 38_40_en1 | 38 | 40 | 1 | 6 | 51 | 57 | 89,47 | 0 | 0 | 0 | 0 | 0 | 0 |
| 143 | 1x1 | 38_40_min_EN | 38_40_en1 | 38 | 40 | 1 | 22 | 55 | 77 | 71,43 | 0 | 0 | 0 | 0 | 0 | 0 |
| 144 | 1x1 | 38_40_min_EN | 38_40_en1 | 38 | 40 | 1 | 16 | 22 | 38 | 57,89 | 0 | 0 | 0 | 0 | 0 | 0 |
| 145 | 1x1 | 38_40_min_EN | 38_40_en1 | 38 | 40 | 1 | 12 | 25 | 37 | 67,57 | 0 | 0 | 0 | 0 | 0 | 0 |
| 146 | 1x2 | 38_40_min_EN | 38_40_en1 | 38 | 40 | 1 | 16 | 100 | 116 | 86,21 | 25 | 25 | 0 | 0 | 0 | 0 |
| 147 | 1x2 | 38_40_min_EN | 38_40_en1 | 38 | 40 | 1 | 24 | 35 | 59 | 59,32 | 5 | 14,29 | 0 | 0 | 0 | 0 |
| 148 | 1x2 | 38_40_min_EN | 38_40_en1 | 38 | 40 | 1 | 49 | 36 | 85 | 42,35 | 2 | 5,56 | 0 | 0 | 0 | 0 |
| 149 | 1x2 | 38_40_min_EN | 38_40_en1 | 38 | 40 | 1 | 17 | 15 | 32 | 46,88 | 0 | 0 | 0 | 0 | 0 | 0 |
| 150 | 1x2 | 38_40_min_EN | 38_40_en1 | 38 | 40 | 1 | 23 | 29 | 52 | 55,77 | 3 | 10,34 | 0 | 0 | 0 | 0 |
| 151 | 2x1 | 38_40_min_EN | 38_40_en1 | 38 | 40 | 1 | 8 | 60 | 68 | 88,24 | 41 | 68,33 | 8 | 27 | 22,86 | 77,14 |
| 152 | 2x1 | 38_40_min_EN | 38_40_en1 | 38 | 40 | 1 | 4 | 39 | 43 | 90,7 | 20 | 51,28 | 8 | 10 | 44,44 | 55,56 |
| 153 | 2x1 | 38_40_min_EN | 38_40_en1 | 38 | 40 | 1 | 5 | 82 | 87 | 94,25 | 19 | 23,17 | 12 | 6 | 66,67 | 33,33 |
| 154 | 2x1 | 38_40_min_EN | 38_40_en1 | 38 | 40 | 1 | 4 | 42 | 46 | 91,3 | 28 | 66,67 | 11 | 15 | 42,31 | 57,69 |
| 155 | 2x1 | 38_40_min_EN | 38_40_en1 | 38 | 40 | 1 | 19 | 26 | 45 | 57,78 | 35 | 134,62 | 15 | 18 | 45,45 | 54,55 |

| | | | | | | | | | | | | | | | | |
|---|---|---|---|---|---|---|---|---|---|---|---|---|---|---|---|---|
| 156 | 2x2 | 38_40_min_EN | 38_40_en1 | 38 | 40 | 1 | 23 | 61 | 84 | 72,62 | 12 | 19,67 | 4 | 8 | 33,33 | 66,67 |
| 157 | 2x2 | 38_40_min_EN | 38_40_en1 | 38 | 40 | 1 | 17 | 37 | 54 | 68,52 | 11 | 29,73 | 1 | 6 | 14,29 | 85,71 |
| 158 | 2x2 | 38_40_min_EN | 38_40_en1 | 38 | 40 | 1 | 6 | 25 | 31 | 80,65 | 9 | 36 | 4 | 5 | 44,44 | 55,56 |
| 159 | 2x2 | 38_40_min_EN | 38_40_en1 | 38 | 40 | 1 | 0 | 20 | 20 | 100 | 12 | 60 | 3 | 7 | 30 | 70 |
| 160 | 2x2 | 38_40_min_EN | 38_40_en1 | 38 | 40 | 1 | 11 | 47 | 58 | 81,03 | 0 | 0 | 0 | 0 | 0 | 0 |
| 161 | 1x1 | 38_50_min_EN | 38_50_en1 | 38 | 50 | 1 | 7 | 47 | 54 | 87,04 | 14 | 29,79 | 3 | 3 | 50 | 50 |
| 162 | 1x1 | 38_50_min_EN | 38_50_en1 | 38 | 50 | 1 | 27 | 60 | 87 | 68,97 | 42 | 70 | 31 | 7 | 81,58 | 18,42 |
| 163 | 1x1 | 38_50_min_EN | 38_50_en1 | 38 | 50 | 1 | 9 | 26 | 35 | 74,29 | 6 | 23,08 | 5 | 1 | 83,33 | 16,67 |
| 164 | 1x1 | 38_50_min_EN | 38_50_en1 | 38 | 50 | 1 | 4 | 50 | 54 | 92,59 | 39 | 78 | 24 | 5 | 82,76 | 17,24 |
| 165 | 1x1 | 38_50_min_EN | 38_50_en1 | 38 | 50 | 1 | 17 | 35 | 52 | 67,31 | 6 | 17,14 | 5 | 0 | 100 | 0 |
| 166 | 1x2 | 38_50_min_EN | 38_50_en1 | 38 | 50 | 1 | 40 | 51 | 91 | 56,04 | 1 | 1,96 | 1 | 0 | 100 | 0 |
| 167 | 1x2 | 38_50_min_EN | 38_50_en1 | 38 | 50 | 1 | 30 | 48 | 78 | 61,54 | 1 | 2,08 | 1 | 0 | 100 | 0 |
| 168 | 1x2 | 38_50_min_EN | 38_50_en1 | 38 | 50 | 1 | 13 | 13 | 26 | 50 | 0 | 0 | 0 | 0 | 0 | 0 |
| 169 | 1x2 | 38_50_min_EN | 38_50_en1 | 38 | 50 | 1 | 29 | 22 | 51 | 43,14 | 1 | 4,55 | 1 | 0 | 100 | 0 |
| 170 | 1x2 | 38_50_min_EN | 38_50_en1 | 38 | 50 | 1 | 36 | 14 | 50 | 28 | 1 | 7,14 | 1 | 0 | 100 | 0 |
| 171 | 2x1 | 38_50_min_EN | 38_50_en1 | 38 | 50 | 1 | 6 | 74 | 80 | 92,5 | 30 | 40,54 | 18 | 5 | 78,26 | 21,74 |
| 172 | 2x1 | 38_50_min_EN | 38_50_en1 | 38 | 50 | 1 | 5 | 43 | 48 | 89,58 | 29 | 67,44 | 15 | 12 | 55,56 | 44,44 |
| 173 | 2x1 | 38_50_min_EN | 38_50_en1 | 38 | 50 | 1 | 1 | 54 | 55 | 98,18 | 41 | 75,93 | 29 | 7 | 80,56 | 19,44 |
| 174 | 2x1 | 38_50_min_EN | 38_50_en1 | 38 | 50 | 1 | 4 | 31 | 35 | 88,57 | 9 | 29,03 | 6 | 2 | 75 | 25 |
| 175 | 2x1 | 38_50_min_EN | 38_50_en1 | 38 | 50 | 1 | 3 | 60 | 63 | 95,24 | 12 | 20 | 6 | 3 | 66,67 | 33,33 |
| 176 | 2x2 | 38_50_min_EN | 38_50_en1 | 38 | 50 | 1 | 7 | 22 | 29 | 75,86 | 6 | 27,27 | 3 | 2 | 60 | 40 |
| 177 | 2x2 | 38_50_min_EN | 38_50_en1 | 38 | 50 | 1 | 5 | 33 | 38 | 86,84 | 2 | 6,06 | 0 | 2 | 0 | 100 |
| 178 | 2x2 | 38_50_min_EN | 38_50_en1 | 38 | 50 | 1 | 11 | 7 | 18 | 38,89 | 1 | 14,29 | 0 | 1 | 0 | 100 |
| 179 | 2x2 | 38_50_min_EN | 38_50_en1 | 38 | 50 | 1 | 7 | 44 | 51 | 86,27 | 0 | 0 | 0 | 0 | 0 | 0 |
| 180 | 2x2 | 38_50_min_EN | 38_50_en1 | 38 | 50 | 1 | 23 | 13 | 36 | 36,11 | 1 | 7,69 | 0 | 1 | 0 | 100 |
| 181 | 1x1 | 38_30_min_EN | 38_30_en2 | 38 | 30 | 2 | 43 | 45 | 88 | 51,14 | 13 | 28,89 | 2 | 1 | 66,67 | 33,33 |
| 182 | 1x1 | 38_30_min_EN | 38_30_en2 | 38 | 30 | 2 | 38 | 44 | 82 | 53,66 | 30 | 68,18 | 21 | 6 | 77,78 | 22,22 |
| 183 | 1x1 | 38_30_min_EN | 38_30_en2 | 38 | 30 | 2 | 19 | 24 | 43 | 55,81 | 13 | 54,17 | 9 | 4 | 69,23 | 30,77 |
| 184 | 1x1 | 38_30_min_EN | 38_30_en2 | 38 | 30 | 2 | 14 | 26 | 40 | 65 | 14 | 53,85 | 7 | 6 | 53,85 | 46,15 |
| 185 | 1x1 | 38_30_min_EN | 38_30_en2 | 38 | 30 | 2 | 16 | 46 | 62 | 74,19 | 23 | 50 | 14 | 7 | 66,67 | 33,33 |
| 186 | 1x2 | 38_30_min_EN | 38_30_en2 | 38 | 30 | 2 | 43 | 13 | 56 | 23,21 | 1 | 7,69 | 0 | 0 | 0 | 0 |
| 187 | 1x2 | 38_30_min_EN | 38_30_en2 | 38 | 30 | 2 | 45 | 30 | 75 | 40 | 11 | 36,67 | 4 | 4 | 50 | 50 |
| 188 | 1x2 | 38_30_min_EN | 38_30_en2 | 38 | 30 | 2 | 46 | 34 | 80 | 42,5 | 10 | 29,41 | 6 | 4 | 60 | 40 |
| 189 | 1x2 | 38_30_min_EN | 38_30_en2 | 38 | 30 | 2 | 40 | 0 | 40 | 0 | 0 | 0 | 0 | 0 | 0 | 0 |
| 190 | 1x2 | 38_30_min_EN | 38_30_en2 | 38 | 30 | 2 | 13 | 39 | 52 | 75 | 17 | 43,59 | 11 | 5 | 68,75 | 31,25 |
| 191 | 2x1 | 38_30_min_EN | 38_30_en2 | 38 | 30 | 2 | 15 | 38 | 53 | 71,7 | 12 | 31,58 | 0 | 2 | 0 | 100 |
| 192 | 2x1 | 38_30_min_EN | 38_30_en2 | 38 | 30 | 2 | 34 | 33 | 67 | 49,25 | 17 | 51,52 | 5 | 4 | 55,56 | 44,44 |
| 193 | 2x1 | 38_30_min_EN | 38_30_en2 | 38 | 30 | 2 | 11 | 48 | 59 | 81,36 | 0 | 0 | 0 | 0 | 0 | 0 |
| 194 | 2x1 | 38_30_min_EN | 38_30_en2 | 38 | 30 | 2 | 6 | 25 | 31 | 80,65 | 4 | 16 | 0 | 4 | 0 | 100 |
| 195 | 2x1 | 38_30_min_EN | 38_30_en2 | 38 | 30 | 2 | 7 | 39 | 46 | 84,78 | 12 | 30,77 | 0 | 6 | 0 | 100 |
| 196 | 2x2 | 38_30_min_EN | 38_30_en2 | 38 | 30 | 2 | 10 | 41 | 51 | 80,39 | 3 | 7,32 | 2 | 1 | 66,67 | 33,33 |
| 197 | 2x2 | 38_30_min_EN | 38_30_en2 | 38 | 30 | 2 | 8 | 15 | 23 | 65,22 | 6 | 40 | 0 | 3 | 0 | 100 |
| 198 | 2x2 | 38_30_min_EN | 38_30_en2 | 38 | 30 | 2 | 20 | 12 | 32 | 37,5 | 5 | 41,67 | 1 | 4 | 20 | 80 |
| 199 | 2x2 | 38_30_min_EN | 38_30_en2 | 38 | 30 | 2 | 41 | 8 | 49 | 16,33 | 4 | 50 | 1 | 2 | 33,33 | 66,67 |
| 200 | 2x2 | 38_30_min_EN | 38_30_en2 | 38 | 30 | 2 | 39 | 13 | 52 | 25 | 5 | 38,46 | 2 | 0 | 100 | 0 |
| 201 | 1x1 | 38_40_min_EN | 38_40_en2 | 38 | 40 | 2 | 35 | 19 | 54 | 35,19 | 0 | 0 | 0 | 0 | 0 | 0 |
| 202 | 1x1 | 38_40_min_EN | 38_40_en2 | 38 | 40 | 2 | 29 | 36 | 65 | 55,38 | 5 | 13,89 | 3 | 0 | 100 | 0 |
| 203 | 1x1 | 38_40_min_EN | 38_40_en2 | 38 | 40 | 2 | 19 | 28 | 47 | 59,57 | 0 | 0 | 0 | 0 | 0 | 0 |
| 204 | 1x1 | 38_40_min_EN | 38_40_en2 | 38 | 40 | 2 | 23 | 32 | 55 | 58,18 | 3 | 9,38 | 3 | 0 | 100 | 0 |
| 205 | 1x1 | 38_40_min_EN | 38_40_en2 | 38 | 40 | 2 | 35 | 32 | 67 | 47,76 | 3 | 9,38 | 3 | 0 | 100 | 0 |
| 206 | 1x2 | 38_40_min_EN | 38_40_en2 | 38 | 40 | 2 | 14 | 20 | 34 | 58,82 | 1 | 5 | 1 | 0 | 100 | 0 |
| 207 | 1x2 | 38_40_min_EN | 38_40_en2 | 38 | 40 | 2 | 42 | 35 | 77 | 45,45 | 0 | 0 | 0 | 0 | 0 | 0 |

| | | | | | | | | | | | | | | | | |
|---|---|---|---|---|---|---|---|---|---|---|---|---|---|---|---|---|
| 208 | 1x2 | 38_40_min_EN | 38_40_en2 | 38 | 40 | 2 | 21 | 19 | 40 | 47,5 | 1 | 5,26 | 0 | 1 | 0 | 100 |
| 209 | 1x2 | 38_40_min_EN | 38_40_en2 | 38 | 40 | 2 | 23 | 27 | 50 | 54 | 0 | 0 | 0 | 0 | 0 | 0 |
| 210 | 1x2 | 38_40_min_EN | 38_40_en2 | 38 | 40 | 2 | 59 | 56 | 115 | 48,7 | 0 | 0 | 0 | 0 | 0 | 0 |
| 211 | 2x1 | 38_40_min_EN | 38_40_en2 | 38 | 40 | 2 | 39 | 22 | 61 | 36,07 | 0 | 0 | 0 | 0 | 0 | 0 |
| 212 | 2x1 | 38_40_min_EN | 38_40_en2 | 38 | 40 | 2 | 63 | 13 | 76 | 17,11 | 0 | 0 | 0 | 0 | 0 | 0 |
| 213 | 2x1 | 38_40_min_EN | 38_40_en2 | 38 | 40 | 2 | 36 | 13 | 49 | 26,53 | 0 | 0 | 0 | 0 | 0 | 0 |
| 214 | 2x1 | 38_40_min_EN | 38_40_en2 | 38 | 40 | 2 | 25 | 26 | 51 | 50,98 | 0 | 0 | 0 | 0 | 0 | 0 |
| 215 | 2x1 | 38_40_min_EN | 38_40_en2 | 38 | 40 | 2 | 32 | 25 | 57 | 43,86 | 0 | 0 | 0 | 0 | 0 | 0 |
| 216 | 2x2 | 38_40_min_EN | 38_40_en2 | 38 | 40 | 2 | 37 | 23 | 60 | 38,33 | 0 | 0 | 0 | 0 | 0 | 0 |
| 217 | 2x2 | 38_40_min_EN | 38_40_en2 | 38 | 40 | 2 | 20 | 54 | 74 | 72,97 | 1 | 1,85 | 0 | 0 | 0 | 0 |
| 218 | 2x2 | 38_40_min_EN | 38_40_en2 | 38 | 40 | 2 | 17 | 29 | 46 | 63,04 | 0 | 0 | 0 | 0 | 0 | 0 |
| 219 | 2x2 | 38_40_min_EN | 38_40_en2 | 38 | 40 | 2 | 22 | 21 | 43 | 48,84 | 0 | 0 | 0 | 0 | 0 | 0 |
| 220 | 2x2 | 38_40_min_EN | 38_40_en2 | 38 | 40 | 2 | 22 | 31 | 53 | 58,49 | 0 | 0 | 0 | 0 | 0 | 0 |
| 221 | 1x1 | 38_50_min_EN | 38_50_en2 | 38 | 50 | 2 | 15 | 29 | 44 | 65,91 | 0 | 0 | 0 | 0 | 0 | 0 |
| 222 | 1x1 | 38_50_min_EN | 38_50_en2 | 38 | 50 | 2 | 38 | 23 | 61 | 37,7 | 2 | 8,7 | 2 | 0 | 100 | 0 |
| 223 | 1x1 | 38_50_min_EN | 38_50_en2 | 38 | 50 | 2 | 42 | 25 | 67 | 37,31 | 0 | 0 | 0 | 0 | 0 | 0 |
| 224 | 1x1 | 38_50_min_EN | 38_50_en2 | 38 | 50 | 2 | 29 | 8 | 37 | 21,62 | 17 | 212,5 | 9 | 8 | 52,94 | 47,06 |
| 225 | 1x1 | 38_50_min_EN | 38_50_en2 | 38 | 50 | 2 | 33 | 14 | 47 | 29,79 | 0 | 0 | 0 | 0 | 0 | 0 |
| 226 | 1x2 | 38_50_min_EN | 38_50_en2 | 38 | 50 | 2 | 38 | 23 | 61 | 37,7 | 0 | 0 | 0 | 0 | 0 | 0 |
| 227 | 1x2 | 38_50_min_EN | 38_50_en2 | 38 | 50 | 2 | 37 | 20 | 57 | 35,09 | 0 | 0 | 0 | 0 | 0 | 0 |
| 228 | 1x2 | 38_50_min_EN | 38_50_en2 | 38 | 50 | 2 | 36 | 11 | 47 | 23,4 | 0 | 0 | 0 | 0 | 0 | 0 |
| 229 | 1x2 | 38_50_min_EN | 38_50_en2 | 38 | 50 | 2 | 29 | 4 | 33 | 12,12 | 0 | 0 | 0 | 0 | 0 | 0 |
| 230 | 1x2 | 38_50_min_EN | 38_50_en2 | 38 | 50 | 2 | 53 | 30 | 83 | 36,14 | 1 | 3,33 | 0 | 0 | 0 | 0 |
| 231 | 2x1 | 38_50_min_EN | 38_50_en2 | 38 | 50 | 2 | 26 | 27 | 53 | 50,94 | 4 | 14,81 | 2 | 2 | 50 | 50 |
| 232 | 2x1 | 38_50_min_EN | 38_50_en2 | 38 | 50 | 2 | 15 | 12 | 27 | 44,44 | 0 | 0 | 0 | 0 | 0 | 0 |
| 233 | 2x1 | 38_50_min_EN | 38_50_en2 | 38 | 50 | 2 | 12 | 19 | 31 | 61,29 | 5 | 26,32 | 2 | 2 | 50 | 50 |
| 234 | 2x1 | 38_50_min_EN | 38_50_en2 | 38 | 50 | 2 | 12 | 34 | 46 | 73,91 | 3 | 8,82 | 3 | 0 | 100 | 0 |
| 235 | 2x1 | 38_50_min_EN | 38_50_en2 | 38 | 50 | 2 | 15 | 31 | 46 | 67,39 | 6 | 19,35 | 2 | 3 | 40 | 60 |
| 236 | 2x2 | 38_50_min_EN | 38_50_en2 | 38 | 50 | 2 | 6 | 20 | 26 | 76,92 | 5 | 25 | 4 | 1 | 80 | 20 |
| 237 | 2x2 | 38_50_min_EN | 38_50_en2 | 38 | 50 | 2 | 7 | 43 | 50 | 86 | 1 | 2,33 | 0 | 1 | 0 | 100 |
| 238 | 2x2 | 38_50_min_EN | 38_50_en2 | 38 | 50 | 2 | 0 | 12 | 12 | 100 | 2 | 16,67 | 0 | 2 | 0 | 100 |
| 239 | 2x2 | 38_50_min_EN | 38_50_en2 | 38 | 50 | 2 | 18 | 18 | 36 | 50 | 0 | 0 | 0 | 0 | 0 | 0 |
| 240 | 2x2 | 38_50_min_EN | 38_50_en2 | 38 | 50 | 2 | 2 | 9 | 11 | 81,82 | 2 | 22,22 | 1 | 0 | 100 | 0 |
